# Survival Benefits Outweigh Germline Competition Costs in Kin Chimeras

**DOI:** 10.64898/2026.04.22.720247

**Authors:** R Voskoboynik, C Askren, NG Gordon, D Krishnamurthy, AG Larson, JC Yu, M Kowarsky, T Levy, KJ Ishizuka, KJ Palmeri, T Rolander, NF Neff, AM Detwellier, SR Quake, M Prakash, IL Weissman, A Voskoboynik

## Abstract

Chimerism, the coexistence of genetically distinct cells within an organism, is a widespread phenomenon in nature. In the colonial chordate *Botryllus schlosseri*, fusion between compatible colonies can lead to stem cell competition with a single genotype taking over the gonadal and/or somatic tissue of the chimera. Over five years, we quantified the frequency and consequences of chimeric formation among 809 larvae, assessing the rates of fusion after natural settlement and its long-term impact on colony survival. We found larvae preferentially settle near kin, facilitating high rates of successful fusion. Fusion was concentrated around the first month of life, boosting survival during this period of high mortality. At all timepoints chimerism significantly increased survival relative to naive and rejected colonies and extended overall lifespan. However, fusion triggered intense internal competition: in all cases tested, a single genotype achieved dominance over both germline and somatic tissues within several weeks post-fusion, regardless of initial genotype contribution. We estimate that among kin, the survival benefits of chimerism outweigh the potential costs of germline loss, extending Hamilton’s kin selection framework to colonial marine organisms. In this context, chimerism functions as an adaptive strategy that enhances early survival while enabling selection among competing stem cell lineages.

## Introduction

Chimerism, the merging of genetically distinct individuals, is a widespread adaptation observed across nine phyla, including protists, plants, and animals *(1)*. In a discussion on chimerism in the colonial marine chordate *Botryllus*, immunologist Frank Burnet hypothesized chimerism could lead to an “ecological niche” for survival by “parasitism on one’s own kind” *(2,3)*. Sabbadin and Zaniolo (1979) confirmed Burnets prediction using pigment markers, which showed germ cells transfer between fused *Botryllus* colonies, resulting in one genotype “takes over” the entire germline *(4)*. Using colony-specific genetic fingerprints (microsatelite), Pancer et al. (1995), Stoner and Weissman (1996) and Stoner et al. (1999) validated the presence of intraspecific parasitism in *Botryllus*, revealing a “winner/loser” hierarchy between fusion partners *(5-7)*.

Chimerism results in a range of documented outcomes, including one partner’s complete somatic and gonadal takeover, partitioning of soma and germline between partners, and various forms of other chimeric mixtures *(4-8 9,10)*. Transplantation experiments established that these dynamics are mediated by competition among prospective somatic and germline stem cells *(11-13)*. Chimeric formation requires immune suppression, loss of individuality, control over allorecognition, and a potential loss of the genetic line. For this strategy to persist, its benefits must outweigh these substantial costs. However, the fitness benefits and frequency of natural chimeric formation, and the genetic changes created by chimeric formation are poorly characterized (*4-10,14–17*). *Botryllus schlosseri* is one of the closest living invertebrate relatives to vertebrates, able to form chimeras, undergo full body regeneration, and reproduce both sexually and asexually, making it a powerful model for bridging the gap of our understanding on chimerism *(18–23,24,25)*. Colonies originate from sexually produced larvae released in a larval batch from a parent colony. After a short free-swimming period larvae settle in aggregates *(23,26)* and metamorphose into sessile oozoids (colony founders) able to initiate chimeric formation with neighboring colonies through blood vessel fusion (*23-26*) (Fig. 1A-G). Successful fusion occurs if colonies share at least one allele in the highly polymorphic *Botryllus* Histocompatibility Factor (BHF) locus *(23,27)*. Colonies that have successfully formed a chimera will share resources and coexist, with resorption of one chimeric partner potentially occurring post-fusion *(24,28,29)*. Conversely, if no BHF allele is shared, colonies mount an immune incompatibility response characterized by a pigmented Point of Rejection (POR) *(23,24,27,30)* (Fig. 1E). Un-related colonies rarely share BHF alleles making them unable to fuse *(23,27, 25)*. BHF is a genetic check on fusion, ensuring chimeras only form between kin *(23-25)*. Wild *B. schlosseri* oocytes are typically fertilized by sperm derived from multiple colonies, generating larval batches of full and half siblings *(23)*. Following the allelic inheritance of BHF, it’s predicted that paired full-sibling colonies will fuse at a rate of 75%, while half siblings (common from wild-caught colonies) will fuse at a rate of 50% *(23)* (Fig.1H-K). These outcomes align with the monogenetic segregation detailed in the work of Sabbadin, 1962 and Scofield et al. *(23-25)*(Fig.1H,J). Prior work by Grosberg and Quinn *(26)* showed that *B. schlosseri* larvae preferentially settle near kin. This preferential kin settlement of full and half-siblings larvae increases the rate of fusion above the ∼50% fusion rate from randomized pairings *(23)* (Fig.1K). In this paper, we present a long-term study of *B. schlosseri*, tracking colonies from release to death, to quantify rates of chimeric formation and the impact of chimerism on colony survival and genetic composition.

**Fig. 1.**
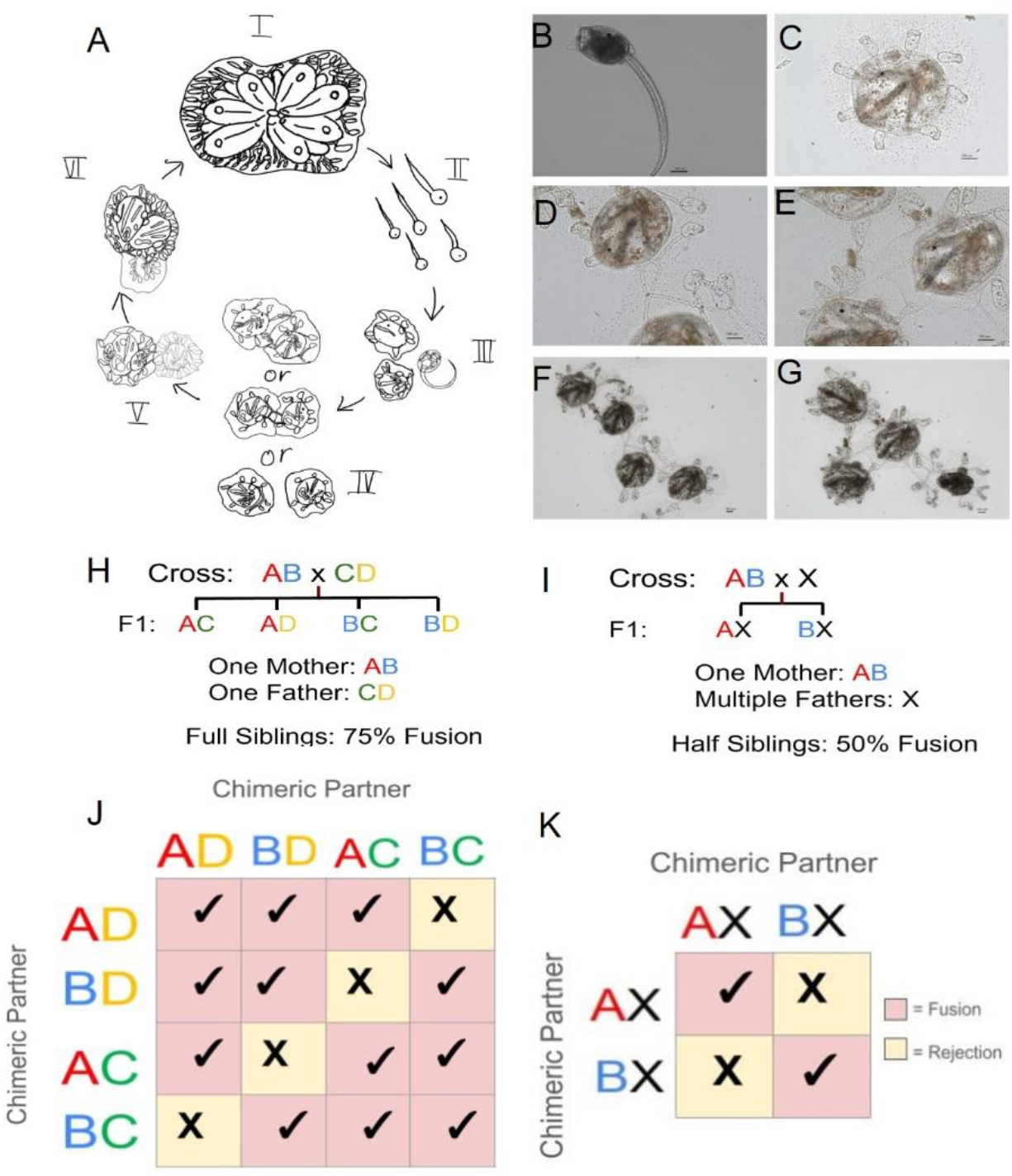
*Botryllus schlosseri* life history and fusion, rejection response. (A) (I) *Botryllus schlosseri* begins life as a sexually produced larva released by a mature parental colony (II) Released larvae have a notochord, dorsal nerve cord, and photoreceptor allowing them to swim and sense light and depth (III) Larvae siblings settle in aggregates, resorbing their tail and other morphological chordate features to metamorphose into a sessile form termed an oozoid (IV) Neighboring oozoids will interact in one of three ways: genetically incompatible oozoids will mount an immune response at the points of contact resulting in highly pigmented Points of Rejection (POR) and no chimeric formation, genetically compatible oozoids may successfully merge blood vessels and form a chimerism to share resources, and alternatively neighboring colonies may avoid interaction remaining adjacent naive individuals. (V) All oozoids undergo weekly full body regeneration following settlement, where the parental oozoid is resorbed by an asexually produced bud that has grown and developed from the oozoid over the course of a week. After the first resorption of a parental oozoid by its asexually produced bud, the structure is termed a zooid, as it has not originated from an oocyte. (I) When more than one bud replaces its parent zooid this creates a multi-zooid structure also called a colony. (B) *In vivo* image of a *Botryllus* larva prior to settlement. (C) A naive oozoid post settlement. (D) Two neighboring oozoids with fused blood vessels. (E) An oozoid, in the top left corner undergoing rejection with two neighboring oozoids. (F) A 3-member multi-chimera, undergoing rejection with a neighboring oozoid (top left corner). (G) An oozoid rejecting the top two members of a 3-member multi-chimera (top left corner), while the 3rd member is being resorbed (bottom right corner). (H) A cross between a single ‘mother’ and ‘father’ colony of AB and CD BHF alleles, and the BHF allelic inheritance of their sexually produced larvae. (I) A visual depiction of the cross between a single “mother” colony and multiple “father” colonies, where none of the fathers share BHF alleles, and the BHF allelic inheritance of their sexually produced larvae. The totality of the fathers who have fertilized the mother colony, contributing to the larval batch, is expressed by ‘X’. (J) Neighboring *Botryllus* colonies require one shared BHF allele to successfully fuse. If no alleles are shared the colonies will reject. All possible BHF alleles of the progeny from a cross of a single “mother” and single “father” are depicted on the sides of a colored table, with the results of the fusion attempts of any two BHF alleles indicated by the color and symbol of the square. A red square with a check mark indicates fusion, while a yellow square with an X mark indicates rejection. Between full siblings there is a 75% predicted fusion rate. (K) All BHF alleles of the progeny from a cross of a single “mother” and multiple “father” colonies are listed on the sides of a colored table, with the color of the correlated square indicating the result of a fusion attempt between progeny with those alleles. Between half-siblings there is a 50% predicted fusion rate.

## Results

### High rate of full-sibling co-settlement following two dimensional water surface interactions

Using a custom made macro-scope developed by the Prakash lab *(31)*, we conducted live image tracking of *B. schlosseri* larvae from the moment of release from a wild type mother colony until settlement and metamorphosis into oozoids (Fig. 2A). While not all released larvae were tracked for the full duration of settlement, a representative group of individuals was successfully tracked. Consistent with earlier observations *(32)* the tracking confirmed classical larval swimming patterns: initial positive phototaxis followed by movement towards the settlement substrate. Through imaging, we identified that larvae demonstrate gravitaxis, swimming upwards towards the surface where they engage in transient interactions with fellow larvae prior to settlement (Fig. 2A). Many organisms face the challenge of locating a specific target, such as a sexual partner (sperm or egg) during spawning or a host in the case of parasites, within a complex environment *(33)*. For marine animals like *B. schlosseri*, the three-dimensional nature of the open water column adds significant complexity to this search *(33)*. Our imaging revealed that larvae swim to the water’s surface prior to interaction, effectively limiting the search for other larvae to two dimensions. This transition reduces the mathematical complexity of locating a target compared to a 3D search. We propose that this 2D search strategy is the primary driver for larval surfacing behavior. We further hypothesize, these surface interactions to be the point where initial kin recognition occurs determining subsequent kin co-settlement.

**Fig. 2.**
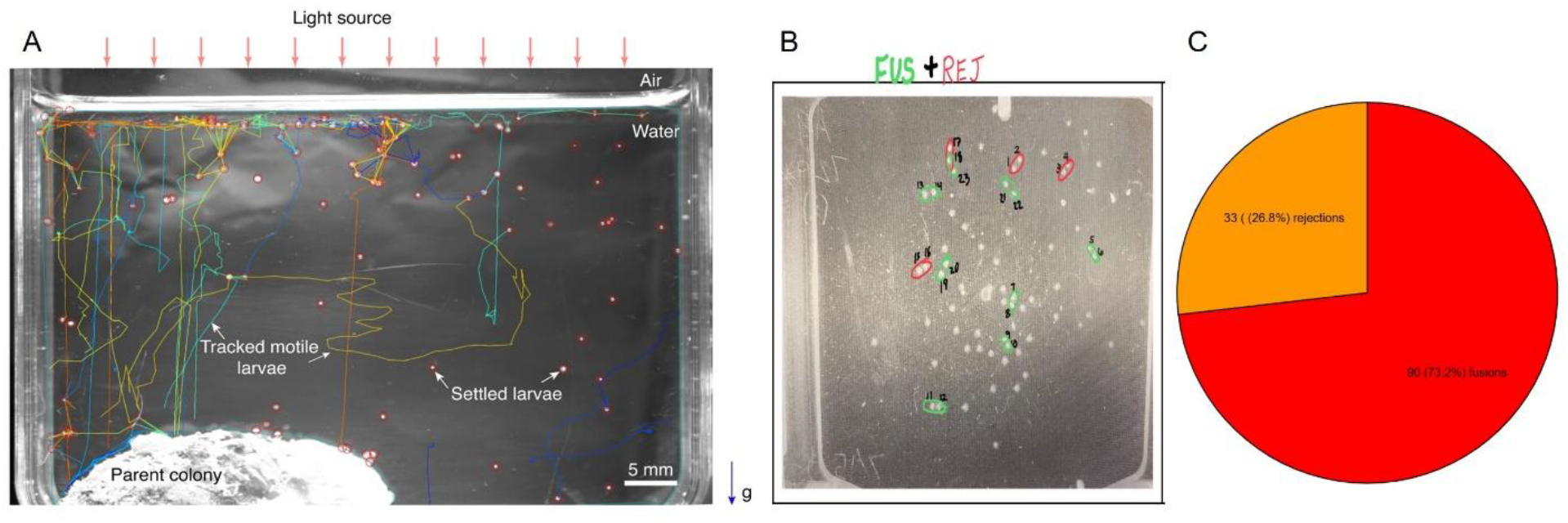
Evidence for full sibling recognition in the larval settlement patterns of *Botryllus schlosseri*. (**A**) Tracking of *Botryllus* larvae siblings from the moment of release from a wild type mother colony, until their settlement. Live imaging and a tracking algorithms were used to follow larvae (**B**) A slide of *Botryllus* oozoids post-settlement 1 day after the larvae had been released by their “mother” colony. Oozoids were tracked and slides were pictured across the experimental period with oozoids that fused by experiment’s end circled in green and those that rejected circled in red. (**C**) A pie chart showing the proportion of fusion (73.2%) to rejection (26.8%) attempt counts in the long and short term experiment trials across all slides. Of the 124 attempts that occurred between two colonies 90 resulted in fusion indicating a shared BHF allele, and 33 resulted in rejection indicating no shared BHF allele between colonies. (Table 1. and DATA S1).

### High rates of successful fusion suggest full-sibling co-settlement

In long-term (5 years) and short-term (2 months) settlement experiments, larvae were tracked from the moment of settlement and metamorphosis until death or the end of the trial. In our observations we followed the interactions between individuals under natural conditions, assessing the frequency of three outcomes: histocompatible fusion (leading to chimera formation), allogeneic rejection, and solitary (naive) development (e.g. Movie S1a and Movie S1b). We counted the total number of contributing genealogies that participated in any histocompatibility reactions. To determine the specific interaction events, we counted each pairwise encounter that resulted in either fusion or rejection. This count included all types of interactions, regardless of whether the participating colonies were naive, already chimeric, or had previously rejected a neighbor (see details in Method section; Table 1; Data S1). In the long-term experiment, 306 individuals were observed. Of these, 59 individuals participated in histocompatibility reactions. These interactions constitute 32 fusion events and 8 rejection events, resulting in a fusion success rate of 80% for the long-term experiment (Table 1; Data S1). Of these 59 interacting individuals, 41 participated in chimerism, 11 were involved in rejection, 4 individuals underwent rejection after forming a chimera and 3 individuals first underwent rejection and then fused with a chimerism (Table 1; Data S1and fig. S2A).

**Table 1.**
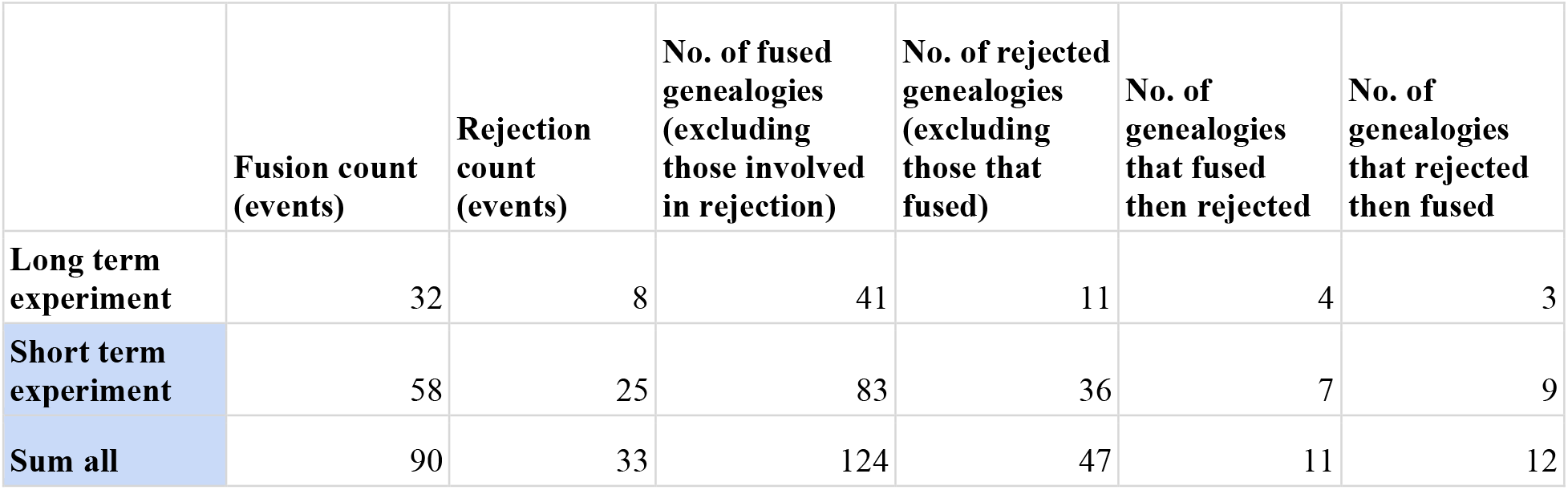
The number of events and distinct genealogies involved in fusion and rejection.

In the short-term experiment, 503 individuals were observed. Of these, 135 individuals participated in histocompatibility reactions. These interactions consisted of 58 fusion events and 25 rejection events, resulting in a fusion success rate of 70% for the short-term experiment (Table 1 and Data S1). A closer look at the 135 interacting individuals will show that 83 participated in chimerism, 36 were involved in rejection, 7 individuals underwent rejection after forming a chimera and 9 individuals formed chimeras after being rejected (Table 1; Data S1 and fig. S2A).

Taken together, 809 individual colonies were observed across both short and long term experiments. Of these, 196 individuals participated in histocompatibility reactions and among them 147 individuals contributed to chimerism at some point (18% of the total observed colonies; Table 1; Data S1). These interactions resulted in a total of 123 fusion/rejection events; with 90 resulting in chimeric formation, and 33 resulting in rejection. This places the rate of successful fusion from all events observed in our experiment at 73% (Fig. 2C). When Scofield et al. *(23)* performed randomized pairing of settled colonies that originated from the same wild type mother, they received a rate of fusion of 50% a number consistent with the rate of fusion expected between half-siblings (Fig. 1I). Our observed numbers from natural settlement are much higher, falling closer to the 75% genetic rate of fusion expected from full-sibling co-settlement. A one-sample proportion z-test comparing our observed fusion rate (73%) to the random settlement expected rate (50%) shows that such an outcome is highly unlikely (z = 5.14, p < 0.001). Comparatively, the expected non-randomized full-sibling co-settlement expected rate (75%) shows no significant difference (z = -0.47, p = 0.61) with our results. (Fig. 1H, Fig. 2B and 2C). These results provide strong evidence that settlement is not random. Instead, larvae are identifying and settling near full-siblings enabling the observed rate of successful fusion.

### Chimeras exhibit significantly enhanced survival, enabling survival during critical early life periods

Our long-term observations revealed a significant survival advantage for chimeric colonies. Of 306 colonies initially monitored, only those that successfully formed chimeras persisted to the conclusion of the 5-year study (Fig. 3A, C, E). Chimeric colonies (n=48) exhibited a median survival time of 273 days, substantially exceeding the 14-day median survival observed in naive colonies (Fig. 3, fig. S1 and Data S1). Conversely, colonies undergoing rejection (n=18) exhibited diminished survival when compared to chimeric colonies. A kaplan-meier curve of these colony types establishes the higher rate of survival that chimeric colonies had over both naive and rejected colonies across the study period (Fig. 3A; Data S1).

**Fig. 3.**
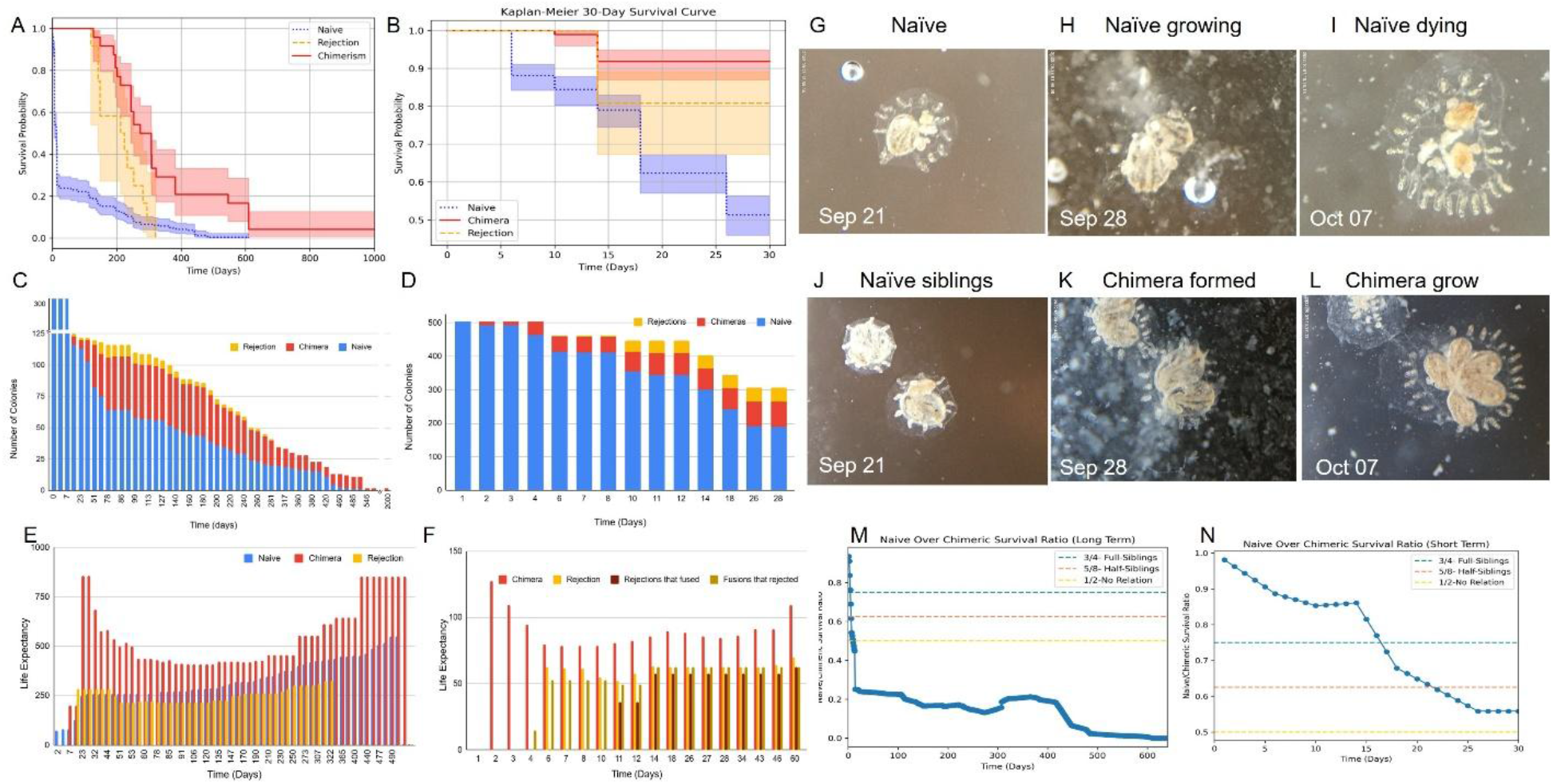
Survival benefits exceed germline competition costs in kin chimeras. (A) A Kaplan-Meier survival curve chart for the naive, chimeric and rejected *Botryllus* colonies in the long-term observational experiment (n=306 colonies). (B) A Kaplan-Meier survival curve for the naive, chimeric and rejected *Botryllus* colonies in the short-term study, during the first 30 days after larvae settlement (n=503 colonies). (C) The total number of naive, chimeric and rejected colonies observed at each time point during the long term study trial. (D) The total number of naive, chimeric and rejected colonies observed at each time point during the short term study trial. (E) Comparison of average life expectancy at each time point among naive, rejecting, and chimeric colonies across the long term observational experiment period with respect to date of rejection and chimeric formation. (F) Comparison of average life expectancy between, rejecting colonies, chimeric colonies, chimeric colonies that later underwent rejection, and rejected colonies that later underwent fusion across the short term trial period (30 days) with respect to date of rejection and chimeric formation. (G-I) An oozoid that remained naive on a settlement slide as observed during the first 26 days following its settlement. Although appearing healthy during the first few weeks, it deteriorated and appeared stressed on day 26, dying shortly thereafter (see also Supplementary Video 1a). J-L) Two oozoids that fused blood vessels and formed a chimera 18 days following settlement. The chimera was observed to be growing and healthy on day 26 (see also Supplementary Video 1b). (M) Calculation of the survival benefit of chimeric formation measured in the long-term observational trial and how it compares to the genetic material lost in chimeric formation based on the degree of relatedness between chimeric partners. The benefits of chimeric formation among kin outweigh the potential germline takeover cost. (N) Calculating the benefits and risks of becoming a chimera based on the degree of relatedness between chimeric partners across the short-term observational experiment. The benefits of chimeric formation among kin outweigh the potential germline takeover cost. Color code: blue-naive, red-chimera, yellow-rejection, brown-rejections that fused, mustard-fusions that rejected. (Table 1 and DATA S1)

In the long term observations, the first 28 days of life constituted the period with the highest rate of die-off: 74% of total naive colonies died during this time period (fig. S1A-B and Data S1). To gain better insight into this period, we conducted a short-term observational experiment (n=503 colonies) focusing on those initial 28 days. This study corroborated our observations, showcasing a significant die-off of 53% of naive colonies during the initial 28 days (Fig. 3B-D). Chimeric formation events were similarly concentrated around this 28 day period (fig. S1C-D and Data S1). The median date of chimeric formation was day six. Chimeric colonies demonstrated significantly enhanced survival compared to naive colonies during the initial 28-day period (p < 0.0001). (Fig. 3A-D; 3G-L; Data S1). Furthermore, comparisons of life expectancy consistently demonstrated a benefit to chimeric colonies. In the long-term observational experiment, we compared average life expectancy among naive, rejecting, and chimeric colonies with respect to the date of rejection or chimeric formation, finding that chimeras exhibited a longer life span at every time point (Fig. 3E). Similarly, in the short-term trial period (30 days), comparison of colonies that were rejecting, chimeric, chimeric then rejected, and rejected then fused also demonstrated a higher life span expectancy for those that formed chimeras (Fig. 3F).

These findings provide robust empirical evidence that chimerism is a highly effective strategy for overcoming early mortality pressures as well as improving animal survival in the long term. While the exact mechanisms for this increased survival (e.g., increased size, shared defense, physiological buffering) were not directly tested here, the outcome is unequivocally positive for the chimeric entity. Conversely, colonies undergoing allogeneic rejection displayed diminished survival compared to chimeras, highlighting the costs associated with failed fusion attempts or prolonged antagonistic interactions (Fig. 3 and fig. S1 and Data S1).

### Inclusive fitness benefits outweigh germline competition costs in kin chimeras

Chimeric formation results in germline competition between participating colonies *(4,6,7-10)*. In many cases, the winner monopolizes the chimera’s reproductive capacity, passing only its own genetic material to the next generation *(4,6,7)* (Movie S2). The losing colony, though contributing to the survival of the chimera, ultimately forfeits its evolutionary fitness by failing to pass on its genes *(4,6,7-10)*. The cost of germline parasitism is mitigated by the genetic relatedness between partners, as the shared proportion of the genome is still transmitted to future generations. We hypothesize that the benefit we have observed chimeras providing in our experiment outweighs the cost incurred through the sacrifice of the germline between closely related colonies. This is consistent with the predictions of Hamilton’s inclusive fitness theory *(34,35)*.

Formation of a chimera is beneficial as long as overall naive survival is less than chimeric survival multiplied by the average proportion of conserved DNA between the two chimeric partners (naive survival < chimeric survival x proportion of conserved DNA). The conserved DNA between two individuals that form a chimera averages to one-half as a baseline and an additional half of whatever proportion of DNA the colonies share. A chimerism between full siblings who share both parents and approximately 50% of their DNA will lead to a germline that contains an average of 75% of the genetic material of the two partners. Meanwhile, a chimerism between two half-siblings which share only one parent and 25% of their DNA, will contain approximately 62.5% of the genetic material of the two partners.

Given that the experiment tracked progeny consisting of both full and half-siblings, these calculations are the most relevant for the trial. Based on these calculations, the benefit of chimeric formation outweighs the cost for full siblings when the ratio of naive to chimeric survival falls below ¾, and for half-siblings when the ratio falls below ⅝. Colonies with no genetic relation experience the greatest loss retaining only 50% of their genetic material, while full-sibling chimeras retain 75% and half-sibling chimeras retain 62.5%.

Our data indicate that naive survival is lower than chimeric survival across all relatedness levels, suggesting that even with the cost of germline loss, the inclusive fitness benefits of chimerism outweigh its risks when partners are sufficiently related. Furthermore, survival analyses (Fig. 3M, 3N) demonstrate that chimeras between more closely related partners will benefit more than those between distantly related or unrelated individuals. These findings support the hypothesis that kin selection plays a role in the evolution of chimerism, favoring its persistence as an adaptive strategy in colonial organisms.

### Multi-Chimerism accelerates resorption

During the study we observed the formation of chimeras between more than two partners. These multi-chimeras formed between three to eight different individual colonies (Fig. 1F-G and fig. S2; Data S1). Proximity played a role in chimeric formation, with chimerism occurring exclusively between the closest settled larvae on the slide. Multi-chimeras similarly occurred among the members of closely settled larvae aggregates.

A common phenomenon within chimerism is resorption, when one individual is resorbed by its fused partner and is no longer physically present on the slide, though its genetic information can live on in the chimeric individual *(6,7)*. According to our data, resorption happened rapidly in multi-chimeras with a larger number of individuals. For example, within a six-individual multi-chimera from our short-term observational slides, it took just six days before one of the individuals began being resorbed, and an additional eight days for another to be resorbed. (fig. S2; Data S1).

### Somatic and germline takeover leading to single genotype dominance in *Botryllus schlosseri* chimeras

To investigate competitive outcomes, seven genetically distinct *B. schlosseri* strains were established. Genetically identical subclones derived from these parent strains were then paired to form five sets of experimental chimeras (Fig. 4A and Data S2). Following fusion, the resorption of one chimeric partner by the other (if occurred) was identified through observed morphological changes. To determine the dominant partner and ascertain the “winner” in terms of somatic and germline contribution, comprehensive genetic analyses were conducted on testes and somatic tissue samples collected from both the chimeras and their naive parents (non-fused) between 1.5 to 5 months post-chimera formation (Data S2). These analyses involved sequencing, alignment of RNA-Seq reads to the *B. schlosseri* genome, and haplotype determination for polymorphic genes (e.g. BHF, Actin2), which were then compared with the alleles of their naive partners (Fig. 4B-C). Since the entire RNA was sequenced and analyzed, we could screen many genes, all of which provided consistent and robust results regarding which genotype took over. This approach established the directionality of resorption and the genetic identity of the winning genotype within each of the five chimera sets, distinguishing the genotypes present in naive versus chimeric somatic (zooids/system) and germline (testes) tissues (Fig. 4A-C and fig. S3.1-3.5 and methods). By combining these allelic fingerprinting results with observations of resorption, individual strains were characterized as “winners” or “losers” within their specific chimeric pairings at each competitive level (resorption, soma, germline; Fig. 4a-c and fig. S3.1-S3.5 and Data S2).

**Fig. 4.**
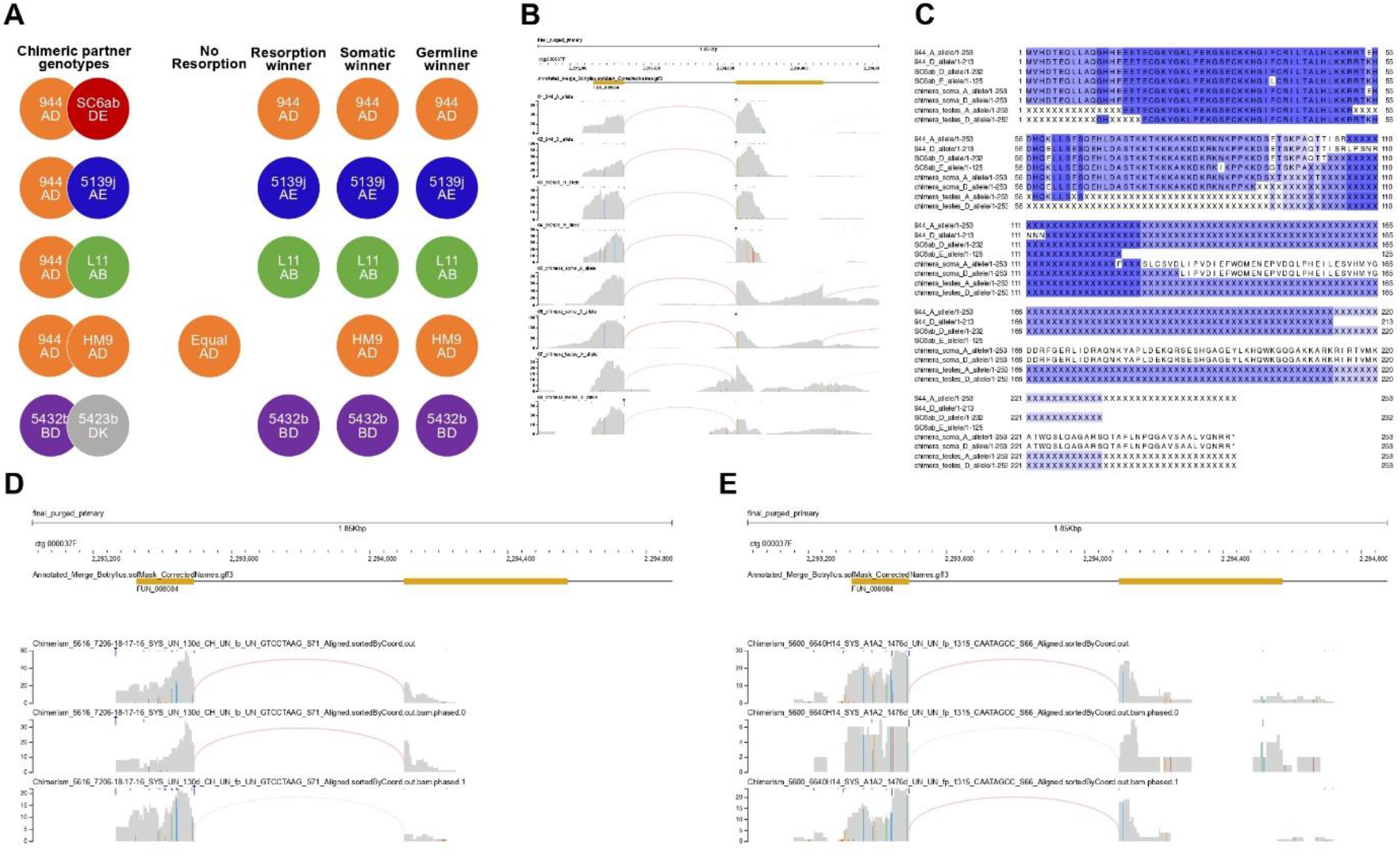
Single genotype dominance in *Botryllus schlosseri* chimeras: somatic and germline takeover 1.5 month to four years following fusion. (A) Directionality of colony resorption and somatic and gametic genetic winners in five *B. schlosseri* chimera sets. The names of chimeric partners and BHF alleles of chimeric partners are included in the illustration. Alignments of RNA-Seq reads to the *Botryllus* genome and haplotype analyses of the BHF and other polymorphic genes were used to identify the specific genotype in naïve and chimeric somatic (zooids/system) and gonadial (test). Samples were collected between 1.5 to 5 months after chimera formation. (B) RNA-Seq read alignments (BHF locus) and haplotype analysis demonstrating takeover by strain 944axBYd196.6.4 (944) (BHF: AD) in a 944-sc6ab chimera (sc6ab BHF alleles: DE) four months post-fusion. Representative naive and chimeric samples are shown. (C) Amino acid alignments of expressed BHF alleles demonstrating takeover by strain 944 (BHF: AD) in a 944-sc6ab chimera (sc6ab BHF alleles: DE) four months post-fusion. Representative naive and chimeric samples are shown. (D) Somatic Takeover 121 days following fusion between 3 siblings on settlement slideOBJ 7206 (individuals 16, 17 & 18), only one genotype is detected (2 BHF alleles; the top alignment shows all read alignments (both alleles); second and third alignments show each allele individually). (E) Single genotype dominance in *Botryllus schlosseri* colony following natural chimera formation (settlement slide 6640, four years (1478 days) post oozoids settlement). 2 BHF alleles; the top alignment shows all read alignments (both alleles); second and third alignments show each allele individually. (fig S3.1-S3.5 and Data S2).

Our investigations revealed that in all five pairs where we successfully sampled both somatic and germline tissue, dominance by a single genotype was established in the chimeras through both somatic and germline takeover. In all the events where a resorption occurred, the resorption winner was also the somatic and germline winner. In the case when both BHF alleles were identical (944 x Hm9; BHF AD for both), a resorption event was not recorded (Fig. 4A marked as “equal”). We investigated competitive outcomes in chimeras derived from the naturally settled oozoids. In a chimera involving three siblings on settlement slide 7206, only a single genotype (indicated by two alleles detected for every polymorphic gene tested) was detected 119 days after their fusion (Fig. 4D). Furthermore, long term dominance was confirmed in a naturally formed chimera on settlement slide 6640, where our haplotyping analyses confirmed a single genotype four years (1478 days) after zooid settlement (Fig. 4E). These findings illustrate a consistent trend towards complete dominance by a single genotype in the long-term outcome of these chimeric interactions.

Next, we took advantage of the ability to form chimeras between colonies with distinct pigment cells to follow their dynamic over 90 days (Movie S2 and Fig. 5). We formed a chimera between an orange colony (dominant pigment) and a blue colony (recessive pigment) isolating it in a dedicated aquarium for time-lapse tracking. Following blood vessel contact and fusion 3 days later (Fig. 5A-C), the former orange partner’s tissues, including the buds, blood vessels, and stem cell niche (endostyle), rapidly adopted a blue phenotype by day 8 (Fig. 5D), and the entire chimera was visibly blue shortly after. This rapid phenotypic shift suggests somatic replacement by the blue genotype. Crucially, all 12 sexually produced oozooids released from the chimera were blue (Fig. 5E-F), despite blue being the recessive inherited color *(4,36-38)*. This result suggests the germline dominance of the blue genotype, whose germline stem cells colonized and took over the reproductive niches of the entire chimeric organism.

**Fig. 5.**
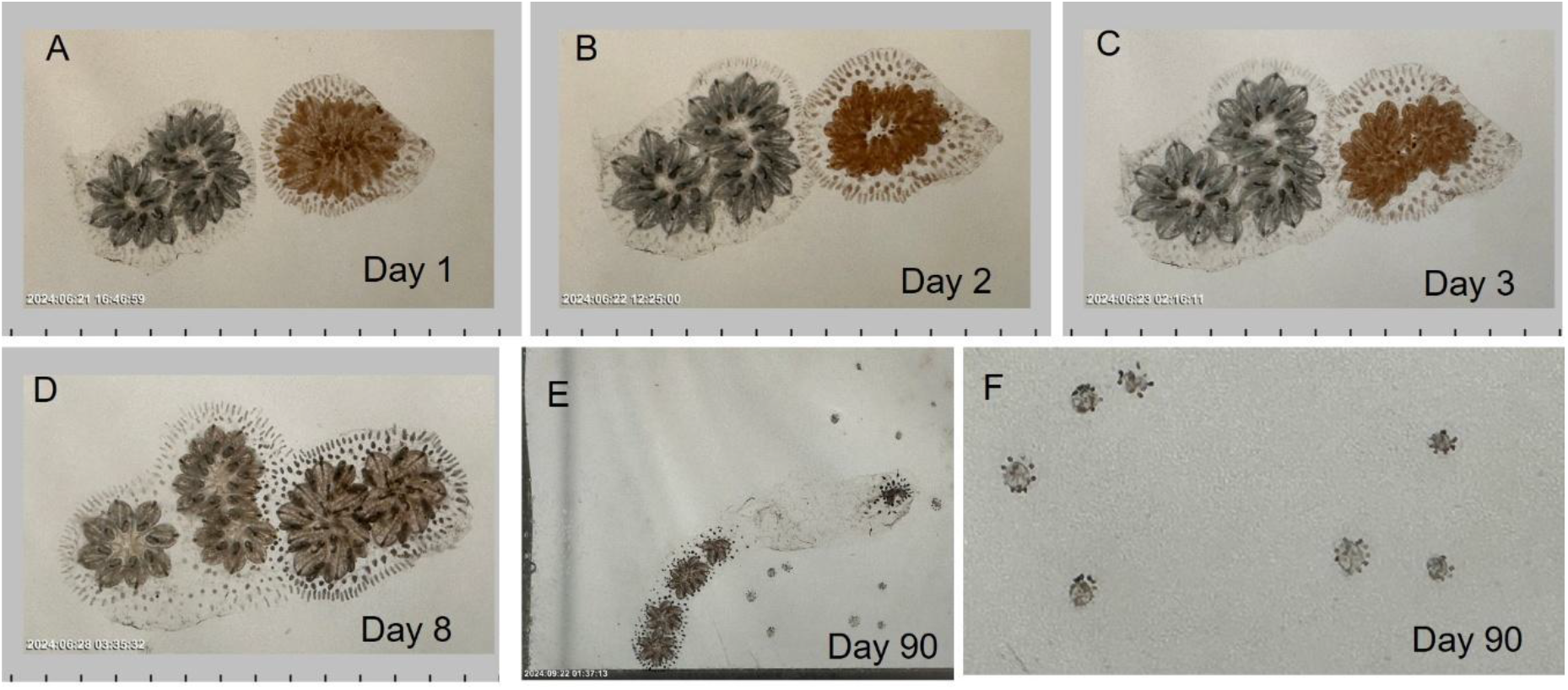
The dynamic of pigment cells in a chimeric partnership of orange and blue individuals across 90 days illustrating both somatic and germline dominance of blue-pigmented cells. **(A)** Two colonies of different genotypes, 7180bX7315a49.1 an orange colony (dominant inheritance) and 7345m a blue colony (recessive inheritance), were placed near each other for an observation and time lapse imaging (day 1). **(B)** The blood vessels (ampullae) of the colonies touched (day 2). **(C)** The colonies’ blood vessels fused, forming a chimera (day 3). **(D)** The blood vessels, buds, and endostyle niche (which harbors adult *Botryllus* stem cells) of both chimeric partners, including the former orange colony (right side) displayed a blue phenotype (day 8). **(E)** Over successive asexual reproduction cycles, the entire chimera adopted a visibly blue phenotype, suggesting a replacement of the orange partner’s somatic tissue. This was observed 90 days following the cut assay setup. Furthermore, all 12 oozooids that settled near the chimera were blue. These oozooids resulted from sexual reproduction between eggs and sperm originating from the chimeric partners (as no other colony was in the tank during the experiment). Since orange is the dominant pigment color compared to blue (based on Mendelian inheritance studies by Sabbadin on *Botryllus* pigment cells^22,36–38^, this suggests that the blue genotype’s germline stem cells (GSCs) actively colonized and dominated the reproductive niches (gonads) of the entire chimera. The two different parts of the chimera maintained separate sexual cycles, which allowed for self-fertilization, yielding offspring whose pigmentation genotype was exclusively blue. This provides further evidence of the transfer and dominance of the blue germline over the orange germline. The full time-lapse imaging followed this chimera for three months, from June 21, 2024, to September 25, 2024 (images taken every five minutes; Movies S2).

### Chimera-Mediated alteration of fusion specificity: an Individual gains compatibility with a previously rejected partner after transient chimerism

Studies on *Botryllus* chimeras, including earlier work by Sabbadin and Astorri (1988) *(24,25)* have demonstrated the principle of “acquired allogeneic tolerance or rejection,” where a colony’s ability to fuse or reject with a partner can change after it forms a chimera with a third, intermediate partner. We also observed that chimera formation influences the success of both fusion and rejection with previously incompatible or compatible partners. In fact, we observed that chimera formation influenced the success of fusion with previously rejected partners on four separate occasions in both our short and long term cases (Data S1 and fig. S2). In the case of settlement slide number 7206 from our short-term study, six individuals near one another fused (fig. S2A). Nearby, a couple of individuals rejected one another. Once those two separated, one of the individuals fused with the multi-chimera for some time before breaking off and heading towards its previously rejected partner again. We found that after having fused with the multi-chimera (and likely receiving its genetic information), that individual then fused with its previously rejected partner. This provides a clear example of how the allele(s) needed to fuse with a previously rejected partner can be obtained through genetic sharing in a chimera. Conversely, chimera formation can also lead to acquired rejection (as shown in Data S1).

Considering that one genotype took over the chimera in all our tested chimeric sets (Fig. 4; fig. S 3.1-3.5), we hypothesized that genotype takeover within a *B. schlosseri* chimera (e.g., where a ‘BC’ genotype becomes dominant in an ‘AB x BC’ fusion) modifies the entire chimera’s functionally expressed BHF allelic profile to primarily reflect that of the dominant ‘BC’ genotype. This acquired ‘BC’ BHF profile subsequently permits the chimera to recognize and successfully fuse with third-party colonies that were previously incompatible with one of the chimera’s original genotype (e.g., a ‘CD’ genotype that rejected the ‘AB’ genotype), because the chimera now shares a common BHF allele (allele ‘C’) with the ‘CD’ genotype, enabling their fusion it will also explain rejection with a colony that before forming a chime(fig. S2B and Data S1). Similarly, this new functional BHF profile can cause the chimera to reject a colony it could have previously fused with, if the new dominant BHF alleles do not match any alleles in the formerly compatible partner.

## Discussion

Our study addresses the central paradox of chimerism in the colonial chordate, *Botryllus schlosseri*: why engage in a strategy that risks total loss of individuality and genetic lineage? Our findings demonstrate that chimerism is not a genetic accident but an essential, highly successful adaptive mechanism that provides a profound survival advantage. This benefit is particularly critical during the first 30 days of life, a period of massive mortality during which 53-74% of naive colonies typically perish. By forming chimeras, colonies dramatically alter their fate, evidenced by the median survival leaping from just 14 days for naive individuals to a remarkable 273 days for fused chimeras. Furthermore, in our 5-year longitudinal study, only chimeric colonies survived to the experiment’s conclusion.

Our data shows that this significant survival advantage is not left to chance but is instead the result of a larval behavior that actively facilitates aggregation with close kin. The high rate of post-settlement colony fusion observed in our study (70-73%) significantly exceeds the 50% baseline expected if settlement/fusion attempts were random among half-siblings *(23,24)*. Instead, this fusion frequency aligns closely with the 75% rate expected for full siblings *(23,24)*. This strongly suggests that the kin-settlement behavior reported by Grosberg and Quinn^26^ is more specific than an indiscriminate aggregation of general kin. We propose that this heightened specificity is enabled by the larval search strategy, where the utilization of a two-dimensional (2D) search pattern at the water’s surface *(33)*, provides a plausible mechanism for larvae to efficiently locate and aggregate with specific kin before committing to final benthic settlement.

While the survival benefit is clear, our results confirm and extend previous findings on germline competition and takeover within *B. schlosseri* chimeras *(4–10)*. Using comprehensive genetic analyses, we demonstrated that single genotype dominance is rapidly established, typically within six weeks post-fusion, across both somatic and germline tissues. The somatic “takeover” was consistent across experimentally formed chimeras and observed in naturally formed multi-individual and long-term chimeras, where a single genotype persisted for up to four years. The resorption winner invariably became the somatic and germline winner, underscoring the competitive nature of these interactions and validating Burnet’s (1971, 1979) concept of “intraspecific parasitism.” *(2,3)*. It is important to note that while the “winner takes all” dynamic was observed in all cases we tested, our sample size is limited. Stoner et al. *(6,7)* and Rinkevich & Yankelevich *(10)*, working with larger sets of chimeras, also found that most interactions resulted in a “winner take all” outcome. However, they identified a few cases where the germline winner did not correspond to the somatic or resorption winner, demonstrating that this split can occur.

Despite the clear cost of potential germline annihilation for one partner, our data support the hypothesis that the survival benefits of chimerism outweigh this cost, particularly among kin. The formula we developed, which incorporates the degree of relatedness, indicates that the observed naive survival rates are consistently low enough for chimerism to be beneficial even when factoring in the transmission of only a portion of a “losing” partner’s genes via inclusive fitness. For full-siblings (retaining an average of 75% of the combined genetic material in the victor’s germline) and even half-siblings (62.5%), the survival advantage conferred by fusing is substantial enough to make chimerism an evolutionary sound strategy. Furthermore, our observation that chimeras between more closely related partners exhibit better long-term survival lends additional support to the role of kin selection in the evolution and maintenance of chimerism.

The core evolutionary question, how a high-stakes gamble like chimerism, involving the potential annihilation of one partner’s germline, can be an adaptive strategy, is resolved by the principle of Inclusive Fitness *(34,35)*. Our data strongly support the hypothesis that the immense survival benefits of chimerism outweigh this potential cost, particularly among kin. In a chimeric fusion, the “loser” colony sacrifices its direct reproductive lineage but ensures the survival and propagation of the genes it shares with the successful “winner,” a classic mechanism of kin selection. Our cost-benefit model confirms that the observed survival advantage against extreme early-life mortality is so substantial that it yields a positive inclusive fitness payoff even when factoring in the transmission of only a portion of the losing partner’s genes. Specifically, the benefit is sufficient for half-siblings (expected genetic retention ≅62.5%) and is maximized by preferential fusion with full siblings (expected genetic retention of ≅75%). This mechanism, where relatedness mediates otherwise costly interactions to enhance survival, is paralleled by other well-documented instances of kin-mediated cooperation across diverse insects and vertebrates species *(34,35,39,40)*.

Collectively, our findings paint a picture of chimerism in *B. schlosseri* as a sophisticated adaptive strategy. It balances the immediate, cooperative benefits of enhanced survival against the ultimate, competitive costs of germline selection. In unpredictable marine environments prone to high early mortality, chimerism likely allows for the rapid selection and propagation of the most robust genotypes or stem cell lineages within a local kin group. In conclusion, our research demonstrates that natural chimerism in *B. schlosseri* is a prevalent and highly adaptive phenomenon driven by kin-selected benefits. Larval behaviors facilitate fusion between relatives, leading to chimeric entities that exhibit significantly enhanced survival, particularly through periods of high environmental stress. While these chimeras are arenas for intense internal competition resulting in single genotype dominance in both soma and germline, the substantial survival advantages conferred by fusion, especially among kin, outweigh the risks of germline parasitism. This complex interplay of cooperation and competition underscores chimerism as a key evolutionary strategy for resilience and adaptation in colonial marine organisms.

## Methods

### Observational study of natural chimera formation and survival

We conducted long-term and short-term observational studies to assess the frequency of natural chimera formation and its impact on colony survival among genetically diverse progeny originating from wild-fertilized *B. schlosseri* colonies. Wild *B. schlosseri* colonies were collected from the Monterey Marina docks on various dates between August 2017 and November 2017 (for the long-term cohort) and July 2022 (for the short-term cohort), specifically selecting areas with low *Botryllus* density. The collected colonies, having been naturally fertilized in the wild, were transported to the Hopkins Marine Station Mariculture facility. Parental colonies were individually tied to 3x5 cm glass slides and placed in slide racks, positioned approximately 10cm opposite clean glass slides within an aquarium. Larvae were allowed to naturally settle onto the clean slides across from the parent colony. This process generated two independent cohorts of naturally settled progeny, which were maintained under previously described mariculture procedures *(21)*.

**The long-term cohort** was generated from seven parental colonies (mother colonies 6585, 6640, 6639, 6636, 6636(2), 6637, 6652) and produced seven settlement slides carrying a total of 306 individual oozoids (colonies). Colonies were tracked with weekly observations for the first month post-settlement, followed by observations every 1-3 weeks for the next 100 days. After the initial 100 days, observations were conducted at least once per month until the death of all colonies or the end of the 5-year study period. During each observation, the colony’s condition, zooid number, and developmental stage were noted, alongside any instances of inter-colony contact, histocompatible fusion (chimerism), or allogeneic rejection (Data S1). Data were used to determine the average lifespan of colonies categorized as naive (solitary), fused, or rejected. We also analyzed the distribution and frequency of chimeric formation, rejection, and death events. The long-term cohort was monitored until the final slide survived to 5 years from initial spawning, at which point samples were collected for molecular analysis (Data S2). Multi-chimeras (fusion between three or more individuals) and instances of acquired allogeneic tolerance (fusion between previously rejecting partners after transient chimera formation with a third party), were recorded.

**The short-term cohort** was established to gain deeper insight into the high-mortality and interaction period observed within the first month of the long-term study. This cohort was derived from five parental colonies (mother colonies 7206, 7219 (1), 7219 (2), 7219 (3), 7226) and generated five settlement slides containing a total of 503 individual oozoids.Observations were taken every three days for the first month. During the first 18 days of life, the total number of colonies, the occurrence of fusion/rejection events, and the state of chimeric colonies were recorded. After the initial 18 days, the observation focus shifted to tracking only existing chimeric and rejected colonies (Data S1). Samples from these slides were taken for molecular analyses (Data S2) as well as time-laps imaging. New fusion, rejection, or contact events involving these groups were noted, while solitary naive colonies were no longer tracked. Observations were continued weekly for the subsequent three months.

#### Statistical comparison to randomness

A **one-sample proportion z-test** was used to compare the observed fusion rate (73%) to the expected fusion rate for random half-sibling settlement (50%) and the expected full-sibling co-settlement rate (75%).

#### Larval Tracking

Larvae swimming were tracked using a custom-built macro-scope as described in *(31)*. A wild-type pregnant mother colony, collected in the Monterey Marina, was placed at the bottom of a cell culture flask (Thermo fisher) filled with seawater. A light was positioned above the flask to mimic natural light conditions at sea. Tracking commenced from the moment of larvae release until final settlement and metamorphosis into oozoid (Fig. 2A). Larval movement, including phototaxis and gravitaxis was analyzed from the video recordings using a tracking algorithm as described in *(31)*.

#### Pictures and Timelapse imaging

A smartphone with an in-house produced phone holder and smartphone time-lapse controller were used to take images and perform timelapse imaging.

Kaplan-Meier survival curve Survival data for naive, rejecting, and chimeric colonies were analyzed using Kaplan-Meier survival curves (Fig. 3A-F; Data S1). Code to generate the curve, and calculate survival relative to sibling relationship can be found here: (Kaplan_meier_short_term.ipynb - Colab), (kaplan-meier - Colab) When generating the total number of colonies that rejected and fused we counted colonies that underwent fusion and then rejection (or the reverse rejected then formed chimerism) as both chimeric colonies and rejecting colonies. We excluded these few events from the figures Kaplan-meier, and the calculations of the impact of fusion and rejection on colony lifespan (Data S1).

#### Germline Competition and Inclusive Fitness Calculations

To evaluate the evolutionary advantage of chimerism in the presence of germline competition, we adapted the principle of Hamilton’s Rule (rB > C) within an Inclusive Fitness framework. This model weighs the survival benefit (B) of fusion against the cost (C) of potential germline annihilation, mediated by the genetic relatedness (r) between the fusing partners *(34,35)*.

##### Defining the adaptive threshold

The formation of a chimera is considered adaptive if the enhanced survival of the chimeric entity, multiplied by the proportion of genes the loser shares with the winner, is greater than the survival of a solitary (naive) colony.

This condition is expressed by the inequality:

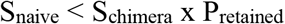

Where: S_naive_: Observed survival rate of naive (solitary) colonies. S_chimera_: Observed survival rate of chimeric colonies. P_retained_: Average proportion of conserved DNA between the two chimeric partners that is successfully transmitted through the winner’s germline.

Calculation of Conserved DNA (P_retained_)

In *Botryllus* chimerism, the germline of the resulting entity is monopolized by a single “winner” genotype. The P_retained_ calculation assumes that the loser loses its direct fitness, but retains indirect fitness proportional to its genetic relatedness to the winner.

The total conserved DNA (P_retained_) is calculated as:

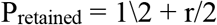

where the 1/2 represents the half of the germline contributed by the winner itself, and r/2 represents the proportion of the loser’s genetic material (r) that is conserved through the winner’s half of the germline.

Based on the known genetic expectations for progeny from wild-fertilized colonies:

Full Siblings (r ≅ 0.50): P_retained_ ≅ 0.50 + 0.50/2 = 0.75 (or 3/4)

Half Siblings (r ≅ 0.25): P_retained_ ≅ 0.50 + 0.25/2 = 0.625 (or 5/8)

Unrelated Colonies (r ≅ 0):P_retained_ ≅ 0.50 + 0 = 0.50 (or 1/2)

##### Determination of Benefit Threshold

By rearranging the adaptive inequality to define a critical survival ratio, we determined the threshold at which the benefit outweighs the cost:

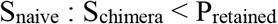

Chimerism is evolutionarily favored when the ratio of naive to chimeric survival is lower than the calculated P_retained_. The observed survival data (S_naive_ and S_chimera_) were then compared to the theoretical P_retained_ values (3/4 for full siblings and 5/8 for half-siblings) to support the role of kin selection.

### Molecular and morphological analysis of dominance

#### Experimental chimera formation

Seven genetically distinct parent strains were established. We subcloned adult colonies and then paired these subclones to generate chimeras (Fig. 4A). The corresponding naive (unpaired) subclones were also maintained for comparison.

The pairing used for chimera formation were:

944axBYd196.6-4 (944) and sc6ab

5423b and 5432b

[5139jL11HMBYSc6ab35]15.9 (5139j) and 944axBYd196.6-4.101 (944)

944axBYd196.6-4 (944) and Hm9axByd196.6.4.219 (Hm9)

944axBYd196.6-4 (944) and L11HM9aBYd196-4-15 (L11).

#### Tissue sampling and RNA sequencing

We performed bulk RNA sequencing (RNAseq) to determine the contributions of individual parent strains to the chimeric tissues. Somatic (zooids/systems stages A-B) and Germline (testes) tissues were collected from lab-paired chimeras and their naive parent 1.5 to 5 months post-fusion (Data S2).We also sampled and sequenced natural formed chimeras including: a chimera formed by a fusion of three individuals, sampled from settlement slides 7206, 119 days following fusion, and five subclones that had undergone natural chimera formation, from settlement slide 6640, four years (1478 days) following oozoids settlement (Data S2). Colonies were frozen in liquid nitrogen and held at -80°C prior to library preparation. RNA was extracted from all tissue samples and sequenced using the methods described in *(21,27)*.

#### Allele calling in chimeras and naive colonies

Reads were aligned to the new *B. schlosseri* reference genome assembly *(41)* (Data S2).Raw reads were trimmed using [trim_galore 0.6.10](https://github.com/FelixKrueger/TrimGalore) set for paired-end reads and a quality setting of 20. The reads were aligned with [STAR 2.7.11b](https://github.com/alexdobin/STAR). Aligned reads were phased and indexed using [samtools 1.19.2](https://github.com/samtools/samtools). Consensus sequences were then obtained using [samtools 1.19.2](https://github.com/samtools/samtools) *(42)* mpileup and [ivar 1.4.2](https://andersen-lab.github.io/ivar/html/manualpage.html) using a script available here (https://github.com/noahggordon/bhf_consensus) to generate consensus sequences from BAM files. To call a base, we required 4x coverage with a minimum of 75% agreement. Nucleic acid consensus sequences were then translated to protein using [EMBOSS Transeq 6.6.0.0](https://launchpad.net/ubuntu/trusty/+package/emboss) We analyzed polymorphic genes (e.g., BHF, FBN2, AHCY1, ACTIN2) to determine the presence of parental alleles in the chimeric and naive tissues (Fig. 4B-C; fig. S3.1-3.5; Data S2). Dominance was established by comparing the allelic fingerprints of the chimeric tissues (soma/germline) to those of the naive partners. Strains were then characterized as “winners” or “losers” at the somatic, and germline levels.Since the entire genome was processed, we could screen many genes, all of which provided consistent and robust results regarding which genotype took over. Resorption, disappearance of one partner’s zooids and buds, was tracked via daily morphological observation.

Visualization and alignment of protein consensuses were done using [Jalview 2.11.4.1](https://www.jalview.org/download/). SNP coverage display was done using [JBrowse 2](https://jbrowse.org/jb2/download/). Figures were created using [inkscape](https://inkscape.org/).

The script for calling BHF alleles is available here: https://github.com/noahggordon/bhf_consensus

### Pictures and Timelapse imaging

A smartphone with an in-house produced phone holder and smartphone time-lapse controller were used to take images and perform timelapse imaging (Video S1, S2A-S2B).

## Acknowledgments

We thank H. Palmeri, C. Patton, L. Hazen. T. Raveh. T. Naik and L. Quinn. for technical advice and help. This manuscript was edited with the assistance of Gemini 3 (Google, version 2025), to improve language style and clarity. The model was used solely as an input for language refinement suggestions and not for data analysis, figure creation, or scientific interpretation. Prompts used included variations of “check grammar, improve flow” applied to draft sections. All edits suggested by Gemini were subsequently thoroughly re-edited by the authors. All scientific content, analyses, and conclusions were originated and verified by the authors.

## Funding

NIH grant R01AG037968 (ILW, SRQ, AV)

NIH grant RO1GM100315 (ILW, SRQ, AV)

NIH grant RO1AG076908 (AV, ILW)

The Chan Zuckerberg investigator program (ILW, AV)

Stinehart-Reed grant (ILW, AV).

Big Ideas for Oceans grant from the Stanford Oceans Department and Stanford Woods Institute for the Environment (AV)

Stanford Bio-X seed grant (AV)

Bio-X undergrad summer fellowship (CA)

Gruss Lipper Postdoctoral Fellowship (TL)

## Author contributions

Conception and design: AV, RV, CA, NGG, KJI, KJP

Mariculture, observation and sample collection: RV, CA, KJI, KJP, AV

Conducting experiments, collecting data: RV, CA, DK, AGL, YCJ, KJI, KJP, TR, TL, AV.

RNA isolation and library preparation: KJP, AV

Sequencing and sequencing analysis: NFN, AM, MK, NGG

Developing a custom microscope and modifying software used for imaging in the research. MP, DK, AGL, TR

Technical support and conceptual advice: AV, NFN, TR, SRQ, MP, ILW

Writing original Draft: RV, CA, AV

Writing review & editing: NGG, TL, KJI, KJP, YCJ, TR, NFN, MP, ILW

Funding acquisition: AV, ILW, SRQ

## Data and Code availability

RNA-seq data have been deposited at the Sequence Read Archive (SRA) under BioProject: PRJNA1289854

The script for calling BHF alleles is available here: https://github.com/noahggordon/bhf_consensus

## Supplemental Figures

**Fig. S1.**
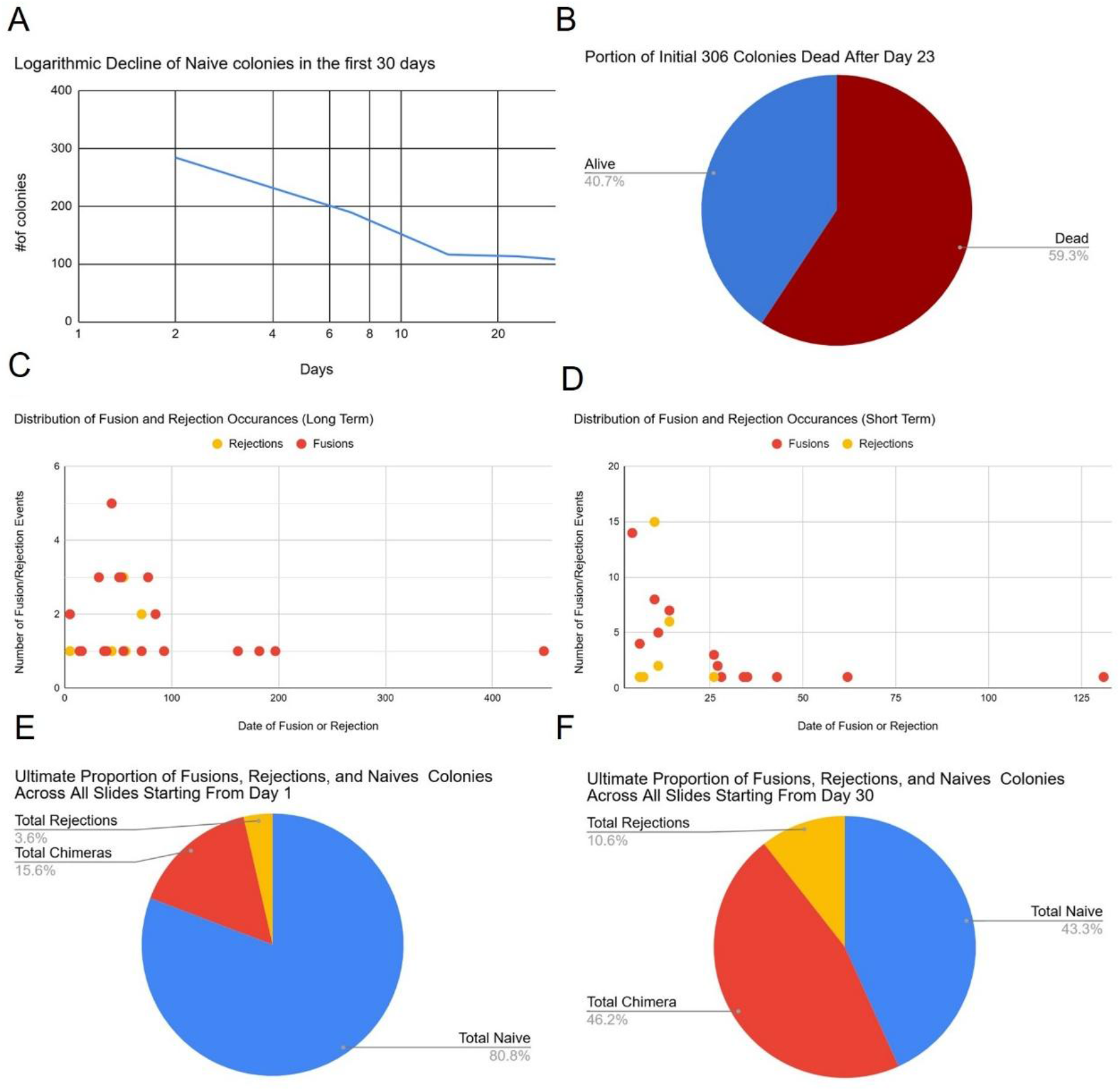
A mass die-off occurred within the first month. Fusion and rejection events were also concentrated during this period. **(A)** A quantification of die off observed in the first 20 days post settlement of the long term experiment, on a logarithmic scale. This graph shows exponential decline during the first 20 days. 30% of initial colonies were dead by week 1and 60% dead by week 2, numbers stayed steady after that point (long term). **(B)** pie chart showing the proportion of initial colonies dead by day 21. **(C-D)** Days during which fusions and rejections occurred. A majority of fusion and rejections events are concentrated around the initial 20 day period (c, long term; d, short term). **(E)** Ultimate proportion of colonies observed across all settlement slides in the long-term experimental period that were chimeras, rejections,or remained naive by experiments end (5 years). **(F)** Ultimate proportion of colonies observed across all settlement slides in the long-term experimental period that were chimeras, rejections, or remained naive colonies by experiments end (5 years) starting from day 30 (excluding the colonies that died in the initial 21 days). (Data S1).

**Fig. S2.**
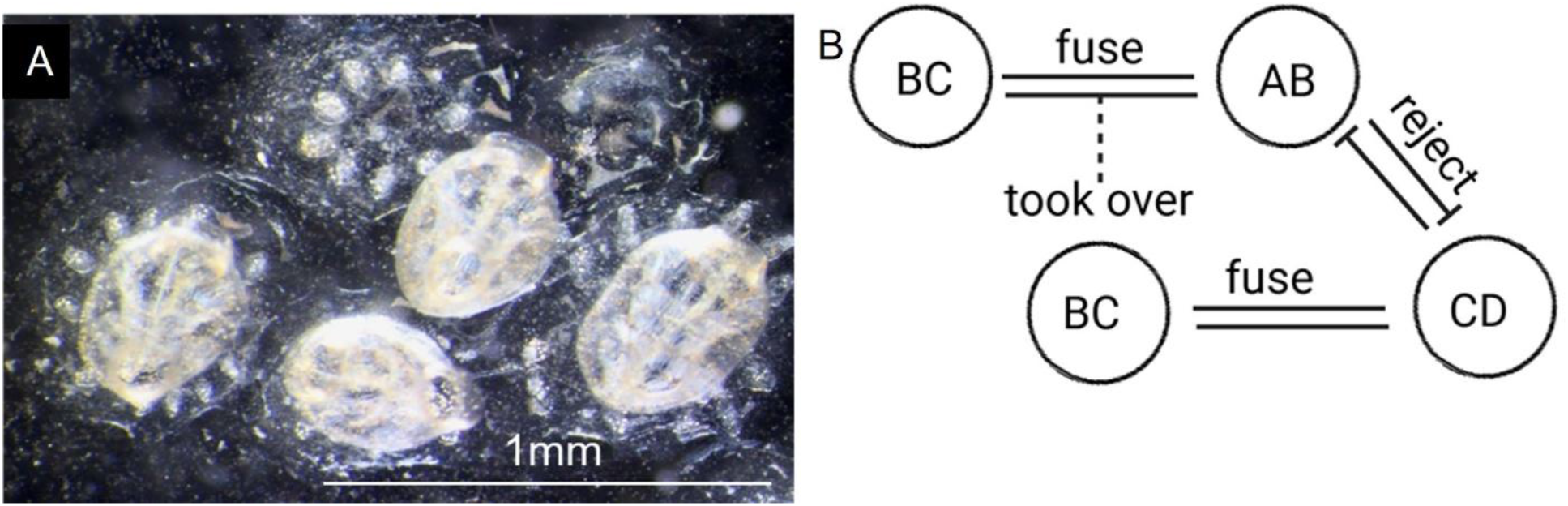
Chimera-mediated alteration of fusion specificity. **(A)** A Six-individual multi-chimera formed by neighboring oozoid siblings. In this chimerism two oozoids initially rejected one another, then after one partner formed a chimera with a separate oozoid it again attempted and succeeded at fusion formation with the original rejected partner.**(B)** Model illustrating how chimera formation between two genotypes that shared a BHF allele (AB x BC) and subsequent genotype takeover modifies BHF allelic profiles (BC) and permits fusion with previously incompatible colonies (CD).

**Fig. S3.1.**
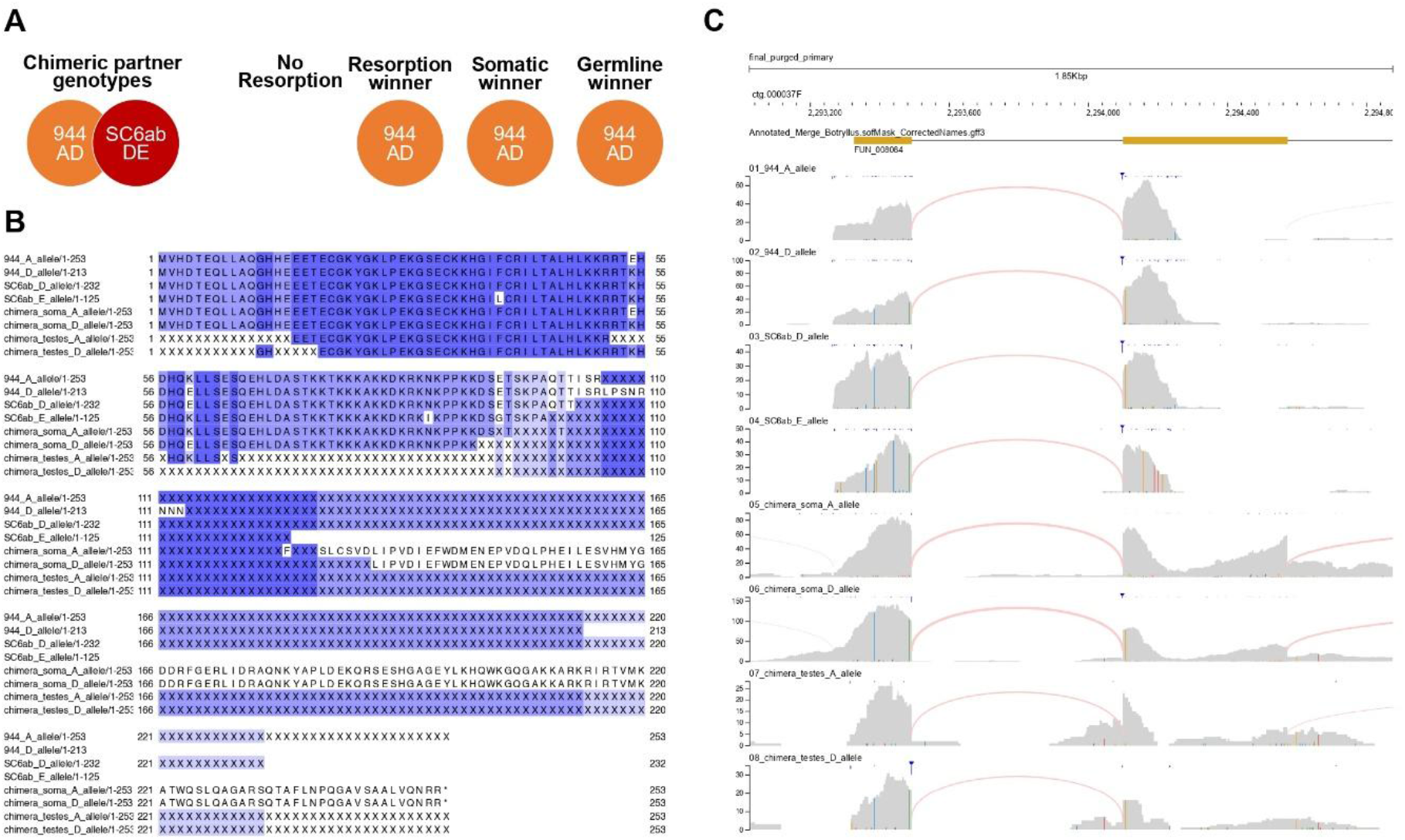
Single genotype dominance in *Botryllus schlosseri* chimeras: somatic and germline takeover four months following fusion. **(A)** Directionality of colony resorption and somatic and gametic identification of the genetic winner in *B. schlosseri* chimera between 944 and Sc6ab. The names of chimeric partners and BHF alleles of chimeric partners are included in the illustration. **(B)** Amino acid alignments of expressed BHF alleles demonstrating takeover by strain 944 (BHF: AD) in a 944-sc6ab chimera (sc6ab BHF alleles: DE) 116 days post-fusion.**(C)** RNA-Seq read alignments (BHF locus) and haplotype analysis demonstrating takeover by strain 944axBYd196.6.4 (944) (BHF: AD) in a 944-sc6ab chimera (sc6ab BHF alleles: DE) 116 days post-fusion. (Data S2).

**Fig. S3.2.**
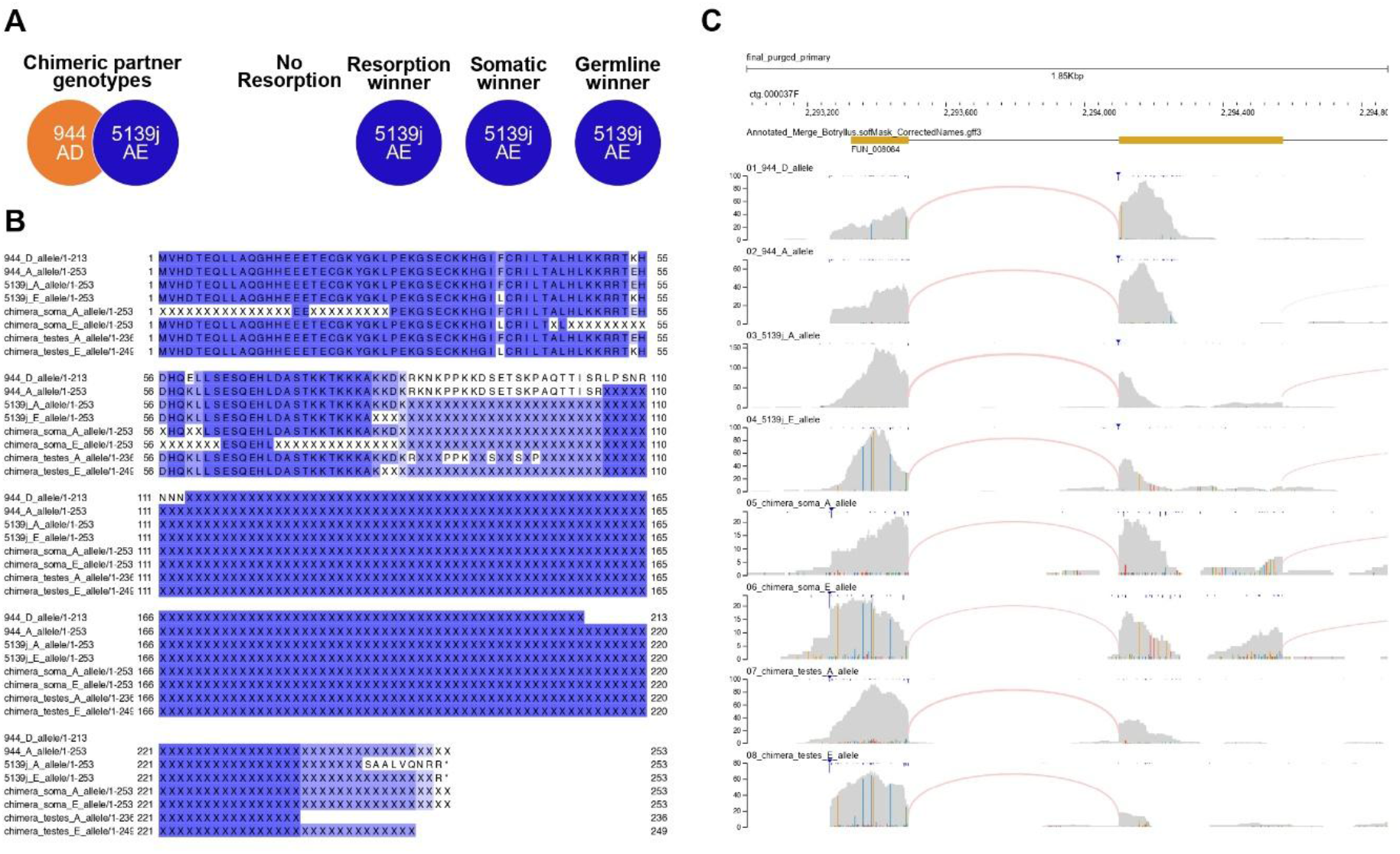
Single genotype dominance in *Botryllus schlosseri* chimeras: somatic and germline takeover two months following fusion. **(A)** Directionality of colony resorption and somatic and gametic identification of the genetic winner in *B. schlosseri* chimera between 944axBYd196.6-4 (944) and [5139jL11HMBYSc6ab35]15.9 (5139j). The names of chimeric partners and BHF alleles of chimeric partners are included in the illustration. **(B)** Amino acid alignments of expressed BHF alleles demonstrating takeover by strain 5139j (BHF; AE) in a 944-5139j chimera 60 days post-fusion. **(C)** RNA-Seq read alignments (BHF locus) and haplotype analysis demonstrating takeover by strain 5139j in a 944-5139j chimera (5139j) (BHF: AE) 60 days post-fusion (Data S2).

**Fig. S3.3.**
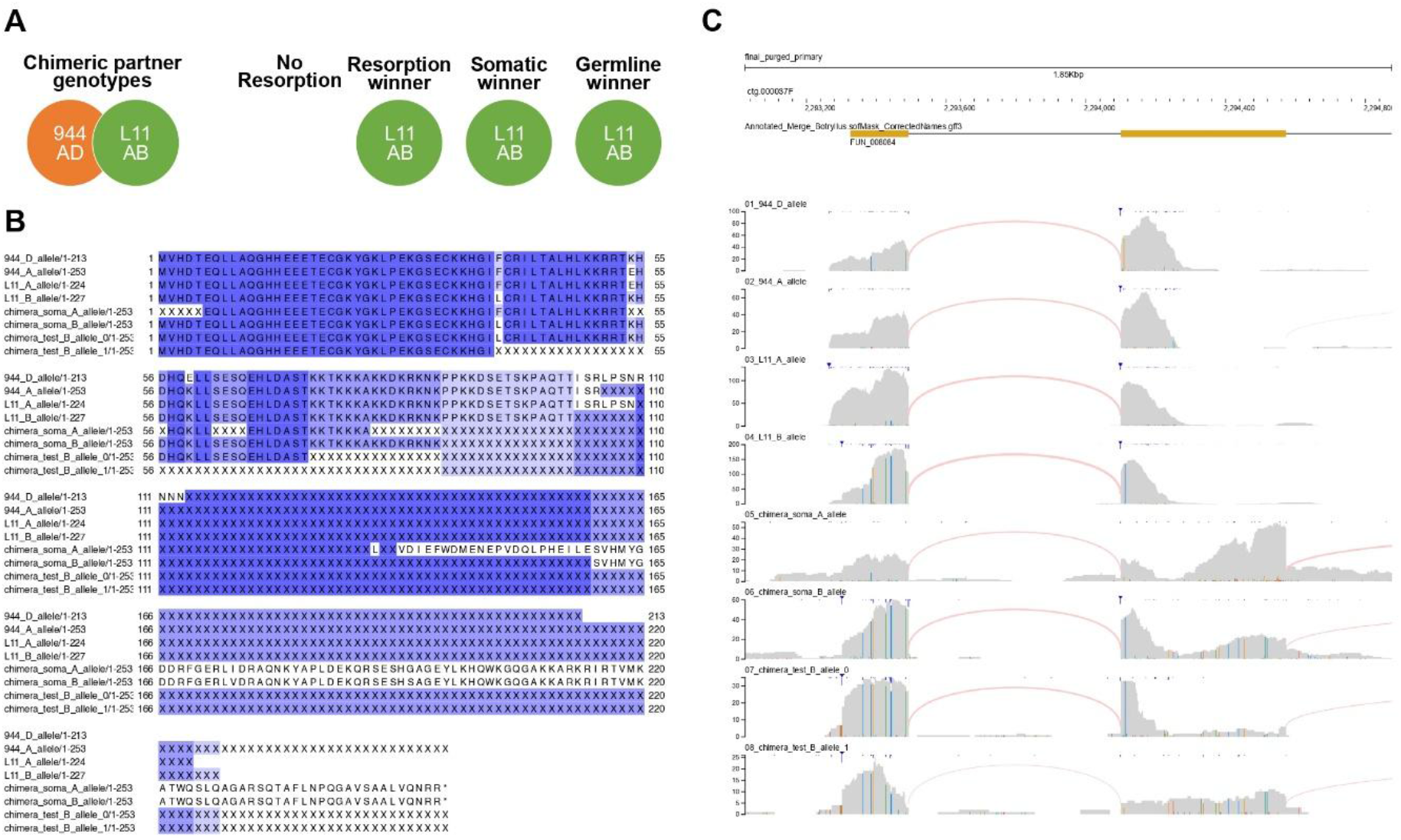
Single genotype dominance in *Botryllus schlosseri* chimeras: somatic and germline takeover 47 days following fusion. **(A)** Directionality of colony resorption and somatic and gametic identification of the genetic winner in *B. schlosseri* chimera between 944axBYd196.6-4 (944) and L11HM9aBYd196-4-15 (L11 called 15 in 2013 paper). The names of chimeric partners and BHF alleles of chimeric partners are included in the illustration. **(B)** Amino acid alignments of expressed BHF alleles demonstrating takeover by strain 5139j (BHF; AE) in a 944-5139j chimera 47 days post-fusion. **(C)** RNA-Seq read alignments (BHF locus) and haplotype analysis demonstrating takeover by strain 5139j in a 944-5139j chimera (5139j) (BHF: AE) 47 days post-fusion (Data S2).

**Fig. S3.4.**
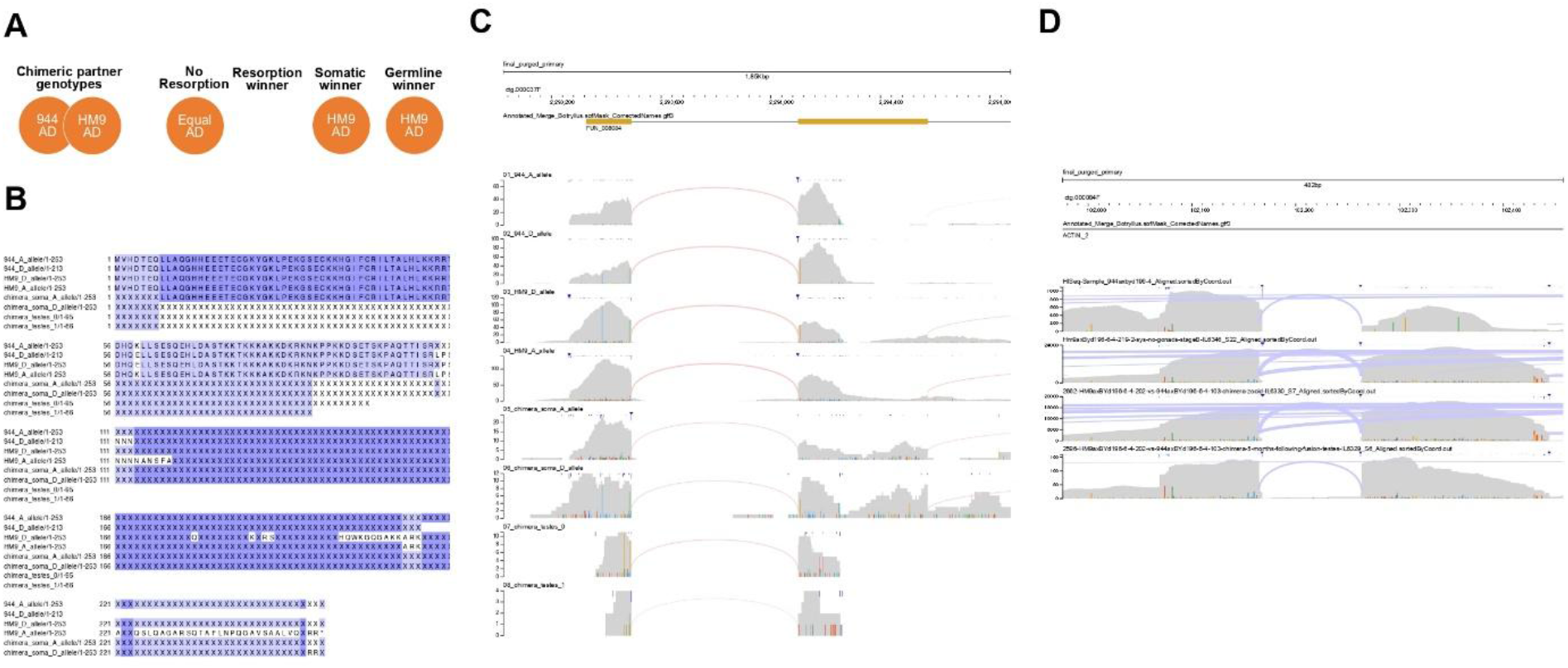
Single genotype dominance in *Botryllus schlosseri* chimeras: somatic and germline takeover 156 days following fusion. **(A)** Directionality of colony resorption and somatic and gametic identification of the genetic winner in *B. schlosseri* chimera between 944axBYd196.6-4 (944) and Hm9axByd196.6.4.219 (Hm9). The names of chimeric partners and BHF alleles of chimeric partners are included in the illustration. In this case where both BHF alleles are identical no resorption winner was observed **(B-C)** Amino acid and RNAseq alignments of expressed BHF alleles demonstrate that both 944 and Hm9 are sharing the same BHF alleles (AD) therefore it is impossible to define a winner genotype based on the BHF locus **(D)** RNA-Seq read alignments to the Actin2 locus demonstrate takeover by strain Hm9 in a 944-Hm9 chimera, 156 days post-fusion both in germline (test) and soma (Data S2).

**Fig. S3.5.**
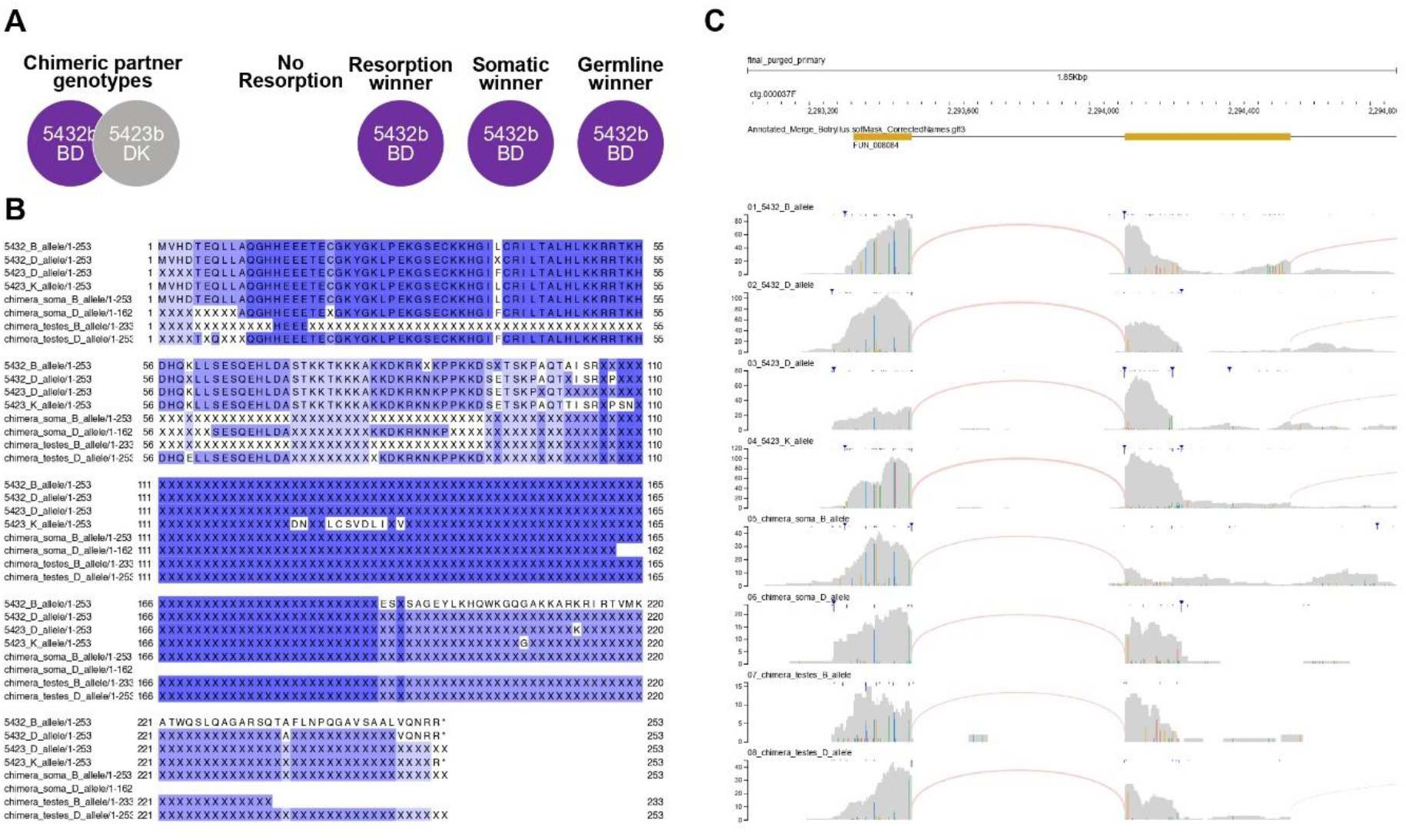
Single genotype dominance in *Botryllus schlosseri* chimeras: somatic and germline takeover 78 days following fusion. **(A)** Directionality of colony resorption and somatic and gametic identification of the genetic winner in *B. schlosseri* chimera between 5423b and 5432b. The names of chimeric partners and BHF alleles of chimeric partners are included in the illustration. **(B-C)** Amino acid and RNAseq alignments of expressed BHF alleles demonstrate that 5432b is the winner genotype based on the BHF alleles (DK) 78 days post-fusion both in germline (test) and soma (Data S2).

**Movie S1A. Early decline and mortality in a naive *Botryllus schlosseri* zooid (Time-Lapse: Days 10-26)**. An oozoid that remained naive on a settlement slide as observed during the first 26 days following its settlement. Although appearing healthy during the first few weeks, it deteriorated and appeared stressed on day 26, dying shortly thereafter. Images were taken every 5 min for 16 days (Sep 21 to Oct 7 2022), starting 10 days after the larvae was settled.

Movie S1B. Successful chimerism and growth of fused *Botryllus schlosseri* zooids (Time-Lapse: Days 10-26; shared the same settlement slide of the zooid observed in Movie S1A) Two oozoids that fused blood vessels and formed a chimera 18 days following settlement. The chimera was observed to be growing and healthy on day 26.

**Movie S2. The dynamics of pigment cells in a chimera between orange and blue individuals over 90 days**, illustrating both somatic and germline dominance of blue-pigmented cells. The full time-lapse imaging followed the chimera formed between 7180bX7315a49.1 (an orange colony) and 7345m (a blue colony). We followed this chimera for three months, from June 21, 2024, to September 25, 2024. Images were taken every five minutes. The date and time are labeled on the upper left side, and the scale bar represents 1 mm.

**Data S1. (separate file)**

**Raw and Summarized Data from *Botryllus schlosseri* Settlement and Allorecognition Experiments:** This data file includes summary tables of all observations from the long-term and short-term settlement slide experiments. Instances of inter-colony contact, histocompatible fusion/chimerism, allogeneic rejection, and date of death are provided. This file contains the tabulated data behind the figures, the raw data used for every graph and analysis presented in the paper is available in titled spreadsheets (e.g., ‘Spreadsheet Fig 3A/M includes the data used to produce these graphs’).

**Data S2. (separate file)**

**Metadata and Genomic Sequencing Metrics for *Botryllus schlosseri* Colonies and Allorecognition Experiments** This table details the genomic sequencing metrics alongside extensive biological and experimental metadata for *Botryllus schlosseri* naive and chimeric colonies analyzed to identify the dominant genotypes within chimeras. To increase confidence in variant calling from *B. schlosseri* predicted genes, only properly paired reads were used for the analyses described herein.

